# The *ATO* gene family governs *Candida albicans* colonisation in the dysbiotic gastrointestinal tract

**DOI:** 10.1101/2025.05.29.656788

**Authors:** Rosana Alves, Faezeh Ghasemi, Wouter Van Genechten, Stefanie Wijnants, Odessa Van Goethem, Cláudia Barata-Antunes, Vitor Fernandes, Patrícia Ataíde, Alexandra Gomes-Gonçalves, Rudy Vergauwen, Qinxi Ma, Ricardo Duarte, Isabel Soares-Silva, Margarida Casal, Alistair J. P. Brown, Patrick Van Dijck, Sandra Paiva

## Abstract

The fungal pathogen *Candida albicans* colonises the human gut where short-chain fatty acids (SCFAs) offer sources of carbon. This fungus harbours one of the largest microbial families of *ATO* (Acetate Transport Ortholog) genes, which encode putative SCFA transport proteins. Here, we generate *C. albicans* null mutants lacking individual or all known putative SCFA transporter genes and compare their phenotypes *in vitro* and *in vivo*. We show that blocking *ATO* function in *C. albicans* impairs SCFA uptake and growth, particularly on acetate. The uptake of acetate is largely dependent on a functional Ato1 (also known as Frp3/Ato3) and it is effectively abolished upon deletion of all *ATO* genes. We further demonstrate that deletion of the entire *ATO* gene family, but not inactivation of *ATO1* alone, compromises the stable colonisation of *C. albicans* in the murine gastrointestinal tract following bacterial disruption by broad-spectrum antibiotics. Our data suggest that the *ATO* gene family has expanded and diversified during the evolution of *C. albicans* to promote the fitness of this fungal commensal during gut colonisation, in part through SCFA utilisation.

**IMPORTANCE:** The human gut is rich in microbial fermentation products such as SCFAs, which serve as key nutrients for both bacteria and fungi. *C. albicans*, a common fungal resident of the gut and a cause of opportunistic infections, carries an unusually large family of *ATO* genes. This study reveals that this *ATO* gene family is required for the efficient uptake of acetate, the most abundant SCFA in the gut, and for stable colonisation of the gut. These findings uncover a new layer of metabolic adaptation in fungal commensals of humans and suggest that transporter gene expansion can shape microbial fitness in response to environmental nutrient signals.

## INTRODUCTION

The mammalian gastrointestinal (GI) tract is colonised by a complex community of microbes. Microbial fermentation of dietary fibres generates short-chain fatty acids (SCFAs), which are known modulators of host-microbe interactions but also essential nutrients that support microbial growth and proliferation in the GI tract (1–3). The efficient utilisation of intestinal SCFAs, either as catabolic or anabolic substrates, is promoted by specific plasma membrane transport systems.

*Candida albicans*, a human fungal pathogen that has evolved both as a commensal of the GI tract and an opportunistic pathogen, encodes transport proteins from two distinct gene families with relevance to SCFA metabolism. The *JEN* family, which includes the carboxylate transporters *JEN1* and *JEN2*, is well characterized and known to mediate carboxylate transport (4, 5). In addition, *C. albicans* has ten genes from the evolutionarily well-conserved acetate uptake transporter (AceTr) family, also known as *ATO* genes, which have been proposed to play a role in SCFA uptake (6). This is the largest known *ATO* family in any microbe (6, 7) and the underlying drivers of the expansion of this family in *C. albicans,* together with the potential functional specialization of Ato proteins in this species, remain obscure. Indeed, acetate is the predominant SCFA in the mammalian GI tract and has long been recognised as a major carbon source for microbes (8, 9).

AceTr members are widely distributed across diverse taxa, including bacteria (SatP/YaaH) (10–12), archaea (AceP) (7, 13, 14), filamentous fungi (AcpA) (15, 16) and yeast (Ady2/Ato1) (6, 7, 17) where they have been reported to play a role in acetate uptake. Acetate transporters from bacterial pathogens of humans are the only AceTr members that have been structurally characterised to date. Two independently solved crystal structures have revealed that bacterial AceTr proteins form a homohexamer complex (Figure 1A) (11, 12). Each of the six proteins consists of six transmembrane segments with their amino- and carboxy-termini located inside the cell (11, 12) (Figure 1B). Structural and biophysical studies initially suggested a channel-like complex (11, 12), but more recent evidence has challenged this idea indicating that acetate translocation might occur via a carrier mechanism (7, 18). Analyses of alpha-fold (19, 20) structures for the *C. albicans* Ato proteins reveal typical architectures analogous to those of bacterial AceTr family members (Figure 1C), suggesting that Ato structural features are evolutionarily conserved from prokaryotic to eukaryotic microbes. This structural conservation supports the idea that the primary function of these membrane proteins as transporters of acetate, and potentially other SCFAs, has been retained in *C. albicans.* However, the *CaATO9* and *CaATO10* genes are most probably nonfunctional. We previously reported that these genes, which encode amino- and carboxy-terminal regions of a full length Ato protein (Figure 1C), appear to have evolved from a single *ATO* gene that has been split by insertion of a transposable element (6). The Ato9 and Ato10 proteins consist of only four and two transmembrane segments, respectively (Figure 1C).

**Figure 1.**
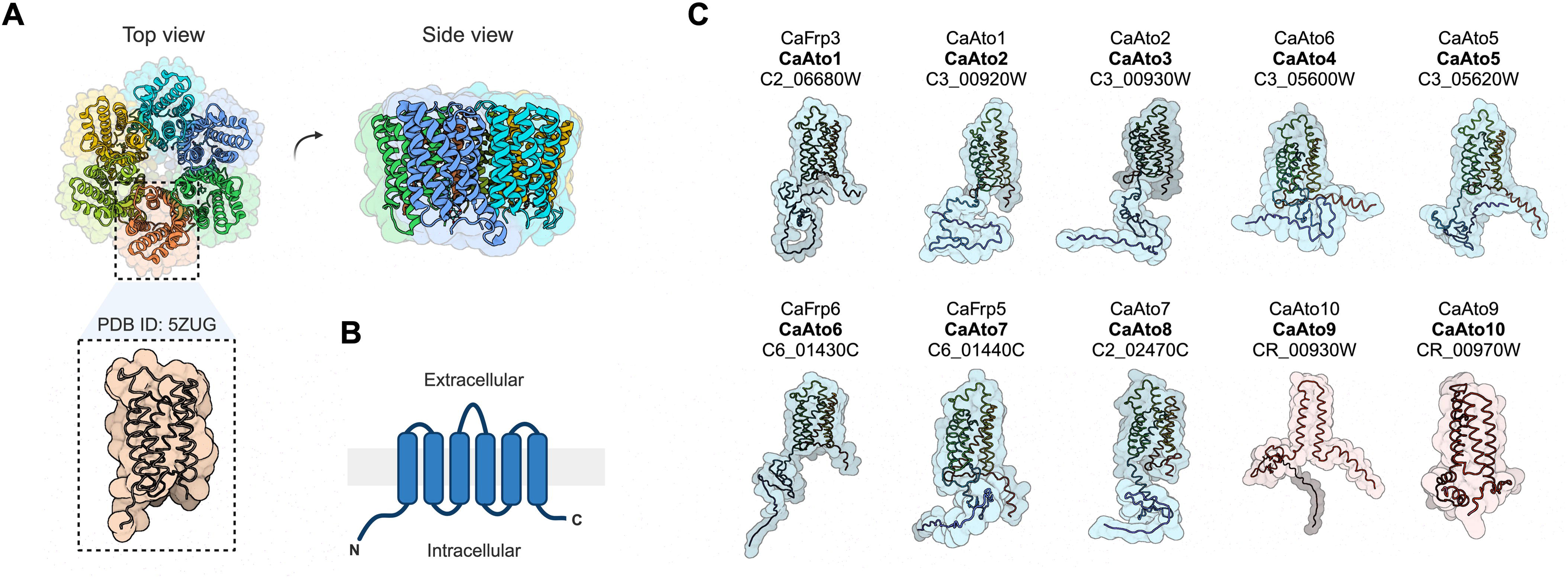
The expanded AceTr transporter family in *C. albicans*. **A.** Top and side views of *Escherichia coli* SatP crystal structure (PDB ID: 5ZUG), a representative prokaryotic member of the acetate uptake transporter (AceTr) family. **B.** Consensus membrane topology for each protomer (11, 12). **C.** Schematic illustration of alpha-fold predictions for the 10 Ato membrane proteins in *C. albicans*. Displayed nomenclature follows new classification (6) (bold) to better reflect and describe their function, while maintaining reasonable consistency with the literature. Standard and systematic names are also included.

It is conceivable that an expansion of the *ATO* gene family might promote rapid adaptation of fungal cells to host associated nutrient gradients thereby offering clear fitness advantages in the context of human colonisation or infection. Supporting this idea, mutations in certain *CaATO* genes impair the fungus’ ability to neutralize the acidic phagolysosome, a trait reported to be linked to hyphal differentiation and survival within macrophages (21, 22). However, it remains unclear whether Ato function is directly associated with virulence (23, 24). Our aim in this study was to investigate the role of the *ATO* gene family in SCFA utilisation and assess its contribution on the stable colonisation of the gut. We provide the first direct evidence, beyond structural inference from bacterial homologs, that Ato proteins mediate acetate transport in *C. albicans*. Moreover, we show that loss of the entire *ATO* gene family impairs stable colonisation of the mammalian gastrointestinal tract, highlighting the collective and environmentally-contingent role of these transporters in fungal commensalism.

## RESULTS

### Inactivation of the ATO gene family in C. albicans

To gain further insights into the function of *C. albicans* Ato proteins, we generated a collection of single and multiple null mutants using CRISPR-Cas9 systems. Additionally, we constructed mutants lacking the monocarboxylate transporter gene *CaJEN1* and/or the dicarboxylate transporter gene *CaJEN2* to serve as controls in our assays (4, 5). Owing to the diploid nature of *C. albicans*, the lack of a traditional sexual cycle (25) and the high degree of genomic plasticity, the generation of marker-free homozygous knockout strains of multigene families remains technically challenging. Nevertheless, we created a set of twelve *C. albicans* mutants, each lacking a specific *JEN* or *ATO* gene, as well as strains with multiple *JEN* and *ATO* knockouts. Furthermore, we successfully eliminated all *JEN* and *ATO* alleles within a single strain.

### ATO disruption abolishes acetate utilisation in C. albicans

We screened these *C. albicans* mutants to reveal differential contributions of the 10 Ato and 2 Jen proteins to acetate uptake (Figure 2A). As acetate transport via specific carriers follows Michaelis-Menten kinetics (26), using a saturating concentration of 4 mM ensured that differences in uptake rates between strains can be attributed to genetic modifications rather than suboptimal substrate availability. Wild-type (WT) control cells displayed 3- to 5-fold higher uptake rates than *ato1*Δ*/*Δ and multiple Ato-deficient cells (Figure 2A, p<0.01). Furthermore, the deletion of *JEN* transporters in combination with *ATO* deletions did not alter the residual acetate uptake capacity (Figure 2A). We then compared the kinetics of radiolabelled acetate transport in WT, *ato1*Δ/Δ, *ato1-10*Δ/Δ, *ato1jen1-2*Δ/Δ and *ato1-10jen1-2*Δ/Δ cells for concentrations between 0.5 mM and 4 mM (Figure 2B). We confirmed that WT control cells displayed a high capacity to transport acetate (*K*_m_ 1.59 ± 0.20 mM and *V*_max_ 1.09 ± 0.06 nmol s^−1^ mg^−1^, Figure 2B), whereas *ato1*Δ/Δ cells showed a significant reduction in acetate uptake (*K*_m_ 4.26 ± 1.42 mM and *V*_max_ 0.51 ± 0.10 nmol s^−1^ mg^−1^, Figure 2B). In the absence of *ATO* genes (*ato1-10*Δ/Δ and *ato1-10jen1-2*Δ/Δ strains), acetate transport was strongly reduced (Figure 2B). The observed residual uptake of radiolabelled acetate in these two strains, which increases proportionally with the tested concentrations of acetate, likely represents passive diffusion of the acid into the cell (Figure 2B).

**Figure 2.**
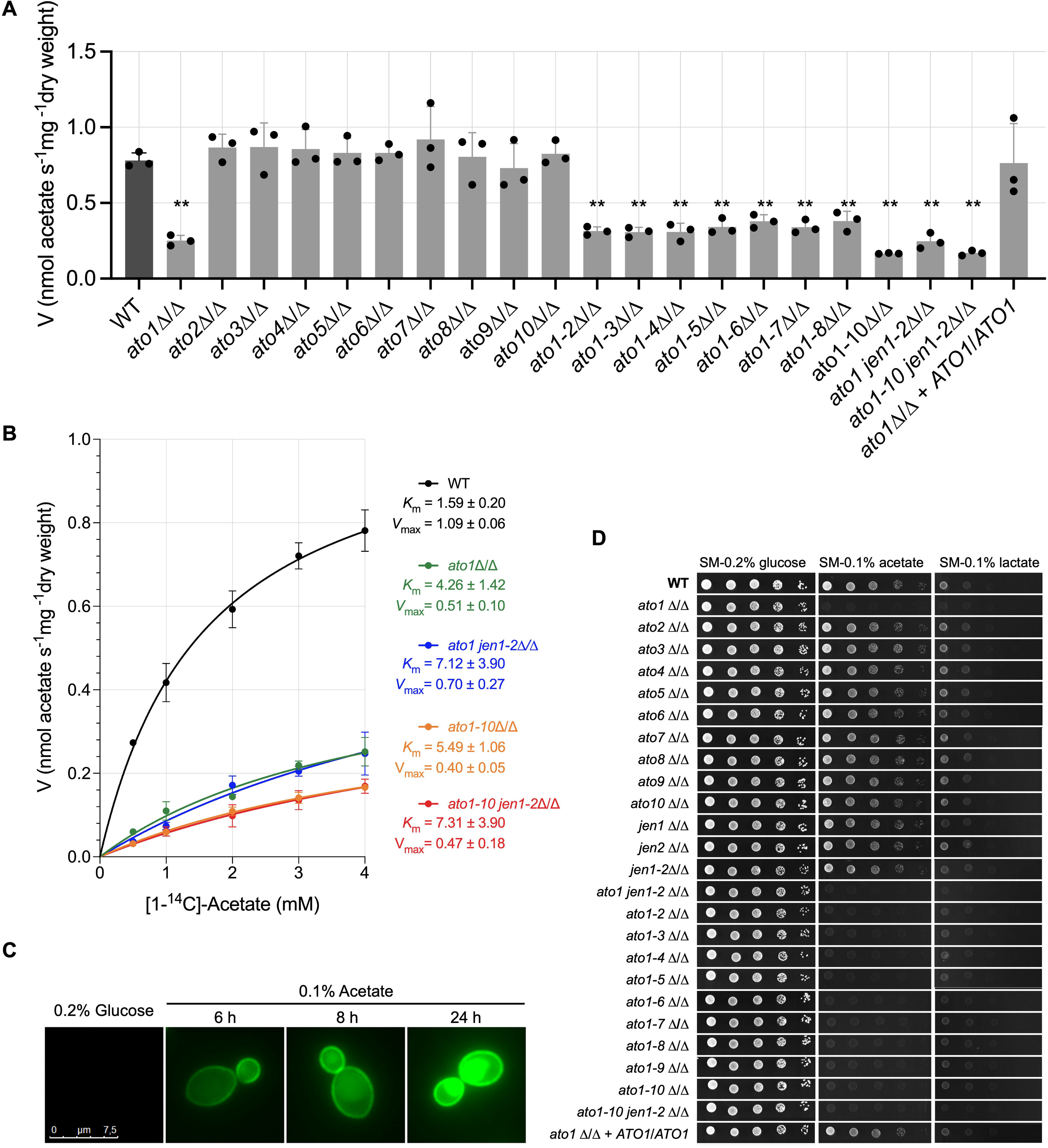
*ATO* disruption in *C. albicans* compromises acetate utilisation. **A.** Uptake of radiolabelled [^14^C]-acetate (4.0 mM, pH 6.0) into *C. albicans* WT and knockout strains grown in Synthetic Complete (SC) medium supplemented with acetate (0.1% v/v, pH 6.0). Error bars indicate mean ± sd of at least 3 independent experiments. Asterisks indicate statistical significance between WT and knockout strains, determined using one-way ANOVA with multiple comparisons tests: **, *p* < 0.01. **B.** Uptake rates of radiolabelled [^14^C]-acetate, at pH 6.0, as a function of acetate concentration, of cells grown in SC medium supplemented with acetate (0.1% v/v, pH 6.0) as sole carbon source. The kinetic parameters (*V*_max_ and *K*_m_) were determined by a computer-assisted nonlinear regression analysis (GraphPad Prism 9.5.0). Error bars indicate mean ± sd of *n=3* independent experiments. **C.** Expression and subcellular localization of Ato1-GFP through time in exponentially growing *C. albicans* cells in Synthetic Minimal (SM) medium supplemented with glucose (0.2% w/v) and then transferred to SC medium supplemented with acetate (0.1% v/v, pH 6.0) as sole carbon source for 6, 8 and 24h. **D**. Growth phenotypes of *C. albicans* WT and knockout strains in SM medium supplemented with specific defined carbon sources: glucose (0.2% w/v), acetate (0.1% v/v, pH 6.0), lactate (0.1% v/v, pH 5.0). Cells were serially diluted, spotted on solid media and incubated at 18 °C for 7 days. Representative data from three independent replicate experiments are shown.

These findings indicate that Ato1 plays an essential role in acetate transport across the *C. albicans* plasma membrane. This hypothesis is supported by Ato1-GFP expression and subcellular localization studies, where a clear Ato1-GFP signal was detected at the plasma membrane when cells were induced in the presence of acetate (Figure 2C). There was a clear temporal increase in *ATO1* expression in response to acetate, in contrast to what is observed in the presence of glucose (Figure 2C). The strong fluorescent signal in the plasma membrane observed at 24 hours was also accompanied by fluorescence within the vacuole. This pattern is consistent with the well-documented intracellular trafficking and recycling of plasma membrane transporters in *C. albicans* and other fungi (27). Internalisation and delivery to the vacuole often occur after prolonged exposure to substrate or as part of turnover and regulation mechanisms (27). Additionally, the growth of *ato1*Δ/Δ cells on acetate as sole carbon source was completely abolished at 18 °C but was restored upon *ATO1* reintroduction (Figure 2D), reinforcing its essential role in acetate utilisation. At this temperature, carboxylic acid uptake via passive diffusion is significantly reduced, making growth on carboxylic acids entirely dependent on the presence of a functional transporter (7). Despite the potential for genetic redundancy across this family, the presence of other *ATO* members in *ato1*Δ/Δ cells did not compensate for the loss of *ATO1* in this growth condition. Indeed, we have recently found that loss of *ATO1* impairs the expression of other *ATO*s in the presence of acetate (28). Furthermore, *ATO1* disruption also impaired cell growth in the presence of lactate, similar to the effect observed for the *JEN1* disruption (Figure 2D). We then evaluated whether the deletion of additional *ATO* genes in *ato1*Δ/Δ cells would confer additive growth defects in the presence of acetate or lactate under a more physiological temperature and growth condition (Figure S1A). Strains carrying additional *ATO* gene deletions, particularly *ato1-10*Δ/Δ and *ato1-10jen1-2*Δ/Δ, did not display any exacerbation of the growth defects observed for *ato1*Δ/Δ cells at 37 °C, either in Synthetic Complete (SC) or Synthetic Minimal (SM) media supplemented with acetate (Figure S1A). Moreover, the growth defects observed in *ato1*Δ/Δ cells were restored upon *ATO1* reintroduction (Figure S1A). As previously observed at 18 °C (Figure 2D), the growth of *ato1*Δ/Δ cells was also impaired in the presence of lactate as the sole carbon source at 37 °C (Figure S1A). However, complete growth abolishment under this condition was only observed upon disruption of *JEN1* (Figure S1A).

### ATO transporter function promotes gut colonisation during antibiotic-induced dysbiosis

Altogether, these observations led us to ask whether both *ATO* and *JEN* families, or *ATO1* alone, influence fungal colonisation in the gastrointestinal tract – a major SCFA-rich ecological niche for *C. albicans*. To test this hypothesis, we performed a series of experiments in a murine model, monitoring the ability of *ato* and *jen* mutants to colonise the gut (Figure 3A). The genotypes of all mutants were confirmed by whole genome sequencing. The resident microbiota was depleted by treating mice with broad-spectrum antibiotics, a prerequisite for *C. albicans* SC5314 colonisation and a well-documented risk factor for candidiasis (29). We compared the bacterial microbiotas of untreated (Figure S2A) versus antibiotic-treated mice (Figures 3B-3D) and the *Candida* infected and uninfected groups by 16S ribosomal RNA gene amplicon sequencing from stool samples (Figures 3B-3D, S2A). Our aim of this amplicon sequencing was to examine the impact of the antibiotic intervention on the intestinal microbiota, and to reveal any potential changes in the gut microbiota associated with colonisation by *C. albicans ato* and *jen* mutants.

**Figure 3.**
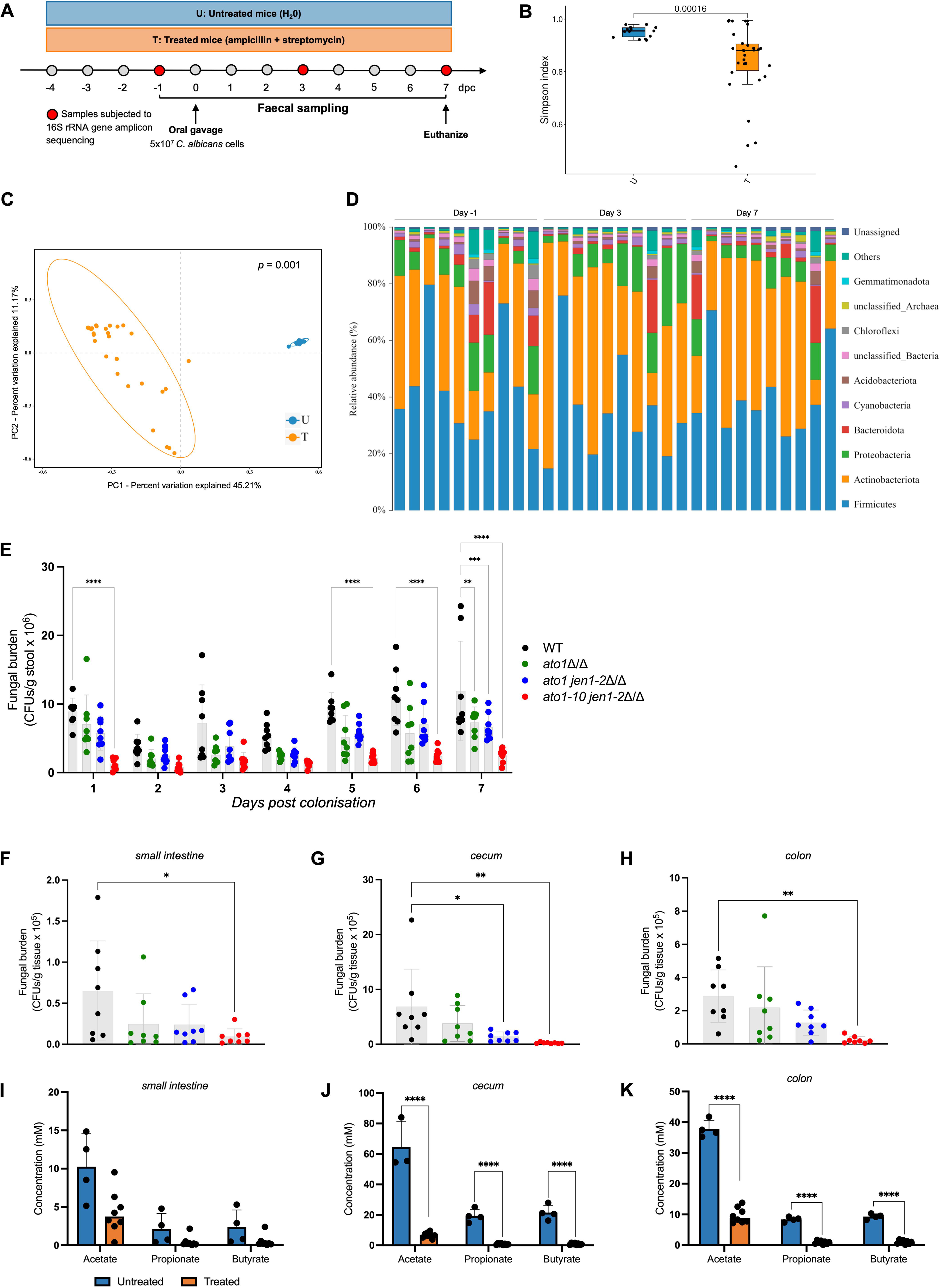
*C. albicans* ATO proteins *allow stable gut occupancy of C. albicans under bacterial dysbiosis.* **A**. Schematic of the experimental design. Mice were colonised with *C. albicans* strains through oral gavage. Experiments were followed for 7 days post colonisation (dpc). **B.** Alpha diversity of untreated and antibiotic-treated groups measured by the Simpson index. Data are shown via interquartile ranges with the median as a black horizontal line. **C.** Beta diversity of untreated and antibiotic-treated groups measured by principal coordinate analysis (PCoA) based on Bray-Curtis distance. **D.** Relative abundance of taxa (%) in antibiotic-treated mice before (day −1) and after *C. albicans* colonisation through oral gavage (day 3 and 7). Each bar represents one independent group of 4 mice. **E**. Fecal colonisation levels (CFUs/g stool) of *C. albicans* strains (*n* = 8 per strain) in antibiotic-treated mice over time. The experiment was performed twice with similar results. Asterisks reflect comparison between strains at individual time points using Tukey’s multiple comparisons test: **, *p* < 0.01; ***, *p* < 0.001; ****, *p* < 0.0001. **F**, **G**, **H**. *C. albicans* levels (CFUs/g tissue) in the small intestine (**F**), cecum (**G**), and colon (**H**) of antibiotic-treated mice (*n* = 8 per strain) at end-point (7 days). Asterisks reflect comparison between strains at individual time points using Tukey’s multiple comparisons test: *, *p* < 0.05; **, *p* < 0.01. **I**, **J**, **K**. *In vivo* quantification of SCFAs measured in the small intestine (**I**), cecum (**J**), and colon colonised by *C. albicans* (**K**) by GC-MS. **, *p* < 0.01; ***, *p* < 0.001; ****, *p* < 0.0001.

The treated mice displayed decreased alpha-diversity compared with untreated controls (*p* = 0.00016, Figure 3B), which was consistent with the antibiotic-induced depletion of bacteria. Similarly, there were significant differences in beta-diversity between untreated and antibiotic-treated samples (*p* = 0.001, Figure 3C). Our data show that the untreated mice were mainly colonised by *Firmicutes* and *Bacteroidota* (Figure S2A). Species of both phyla were the main targets of the administered antibiotics, with *Bacteroidota* species being partially eliminated (Figure 3D). We also observed a significant increase of *Actinobacteriota* and *Proteobacteria* species as result of the antibiotic treatment (Figure 3D).

These phylogenetic alterations in the intestinal microbiome were consistent with a dysbiotic state, often observed after antibiotic treatment (30) and during colonic inflammation (31) in both murine models and clinical studies. *Candida* colonisation did not alter substantially the bacterial microbiota: no significant differences were observed before and after *C. albicans* colonisation (Figures 2D, S2A). Furthermore, our comparisons of groups gavaged with *C. albicans* WT and knockout mutants showed high degrees of similarity in their microbiotas (Figures 3D, S2A).

These analyses were important to ensure that the antibiotic treatment itself did not introduce confounding changes in bacterial community composition, independent of fungal colonisation. However, these groups displayed significant differences in their fungal colonisation levels (Figure 3E). *C. albicans* levels were evaluated by quantifying fungal colony forming units (CFUs) in faeces over the course of seven days. Significantly, none of the tested strains exhibited any fitness defects *in vitro* in the presence of glucose, either in solid (at 18 °C and 37 °C, Figures 2D, S1A) or liquid media (at 30 °C, Figures S1B-S1C).

As expected (32), control mice that were not treated with antibiotics were not stably colonised by *C. albicans* (Figure S2B). In contrast, all tested strains were able to colonise the GI tract of the antibiotic-treated mice (Figure 3E). However, under these conditions, WT cells exhibited increased fitness when compared to *ato1-10jen1*-*2*Δ/Δ cells, with significant differences arising after five days of colonisation (*p* < 0.0001, Figure 3E). Consistent with this result, *ato1-10jen1*-*2*Δ/Δ colonisation levels were also reduced, in comparison to WT, when directly assessed in different gut compartments such as in the small intestine (*p* < 0.01, Figure 3F), cecum (*p* < 0.01, Figure 3G) and colon (*p* < 0.01, Figure 3H). Fungal burdens were consistently lower in the small intestine (Figure 3F) than in the cecum (Figure 3G) or colon (Figure 3H), which is in agreement with previous observations (33). The significant reduction in the colonisation levels observed for the *C. albicans ato1-10jen1-2* mutant compared to both WT and *ato1* strains (Figure 3E-H), suggests that *ATO* gene family members other than *ATO1* promote *C. albicans* fitness during gut colonisation.

We also quantified the levels of the key SCFAs acetate, propionate and butyrate in the different gut compartments by gas chromatography-mass spectrometry (GS-MS) to ensure the physiological relevance of our experimental model. The observed levels of acetate, propionate and butyrate remained aligned with the classical reported ratio of 3:1:1 for these acids in the gut (Figures 3I-K) (33). SCFAs levels were reduced in antibiotic-treated mice, consistent with decreased bacterial fermentation under this condition. Nevertheless, these acetate concentrations remained comparable to those tested and shown to be dependent upon Ato-mediated uptake *in vitro* (Figure 2).

## DISCUSSION

Our study demonstrates that the fungal pathogen *C. albicans* coordinates the assimilation of the key SCFA acetate via the Ato family to facilitate growth and colonisation of the host. Our data are consistent with the idea that AceTr function is retained in *C. albicans* Ato1, with additive effects on acetate utilisation observed as multiple *ATO* genes are eliminated. While inactivation of *ATO1* alone impairs growth on acetate, only the deletion of the entire *ATO* gene family leads to a significant reduction in the gut colonisation levels of *C. albicans*. This suggests functional redundancy and highlights the environmental dependence of *ATO* gene function. The ability to transport acetate confers a metabolic advantage to *Candida* in the GI tract, where this SCFA is abundant and serves as a major available exogenous carbon source. It also enables fungal cells to sustain metabolism and survival when thriving in the diverse and complex niches of the host. The antibiotic-treated murine model essentially reflects the increased risk of candidiasis that antibiotic treatments pose to humans. Indeed, acetate concentrations in the GI tracts of antibiotic-treated mice (Figures 3I-K) closely mirrored those reported in human faeces following broad-spectrum antibiotic therapy (34), thereby validating the physiological relevance of our model.

Antibiotic treatment reshapes the gut environment in multiple, interdependent ways beyond simply reducing SCFA pools. The depletion of commensal anaerobes alters niche availability, nutrient flux, and pH, and can also modulate mucosal immune responses and epithelial barrier function (26–28). Such changes may expose novel adhesion sites and/or relieve colonisation resistance imposed by bacterial competitors. By probing both single- and multi-gene *ato* and *jen* mutants under these complex conditions, our study begins to unravel how SCFA uptake pathways interface with host-microbiota interactions (35). The fact that multiple ATO family members are required for robust colonisation during antibiotic-induced dysbiosis suggests that the functional diversity of this expanded transporter family is critical for fungal persistence in a dynamically changing gut ecosystem.

In conclusion, our findings suggest that, during the evolution of *C. albicans,* the *ATO* gene family has expanded and diversified to promote the fitness of this fungal commensal during gut colonisation, and possibly in other mucosal niches, in part through the efficient assimilation of SCFAs. We propose that ATOs are significant molecular players in the gastrointestinal colonisation program of *C. albicans*. Further studies are required to define how individual Ato proteins are regulated, how they function in concert, and how their activity integrates with broader networks of nutrient sensing and host adaptation.

## MATERIALS AND METHODS

### Materials and data availability

Further information and requests for resources and reagents should be directed to and will be fulfilled by the corresponding author, Sandra Paiva. All unique materials generated in this study are available from the corresponding author with a completed Materials Transfer Agreement. Any additional information required to reanalyse the data reported in this work is also available upon request.

### Strains and culture conditions

All *C. albicans* strains used in this study were derived from the wild-type strain SC5314 and are listed in Supplementary Table 1. Prototrophic strains were routinely cultured in yeast extract-peptone-dextrose (YPD) medium, while nourseothricin-resistant strains were selected on YPD supplemented with 200 µg/mL nourseothricin. For *in vitro* assays, cells were grown at 30 °C, either in Synthetic Minimal (SM: 0.69% w/v YNB from Formedium) or Synthetic Complete (SC: 0.67% w/v YNB from Difco, 0.2% w/v Kaiser SC mixture) liquid media supplemented with specific defined carbon sources: glucose (0.2% or 2% w/v) or acetate (0.1% v/v, pH 6.0). The culture media pH was adjusted with a NaOH solution (12M). Solid media was prepared by adding agar (1.5% w/v) to the respective liquid media.

### Generation of CRISPR-cas9 genome edited *C. albicans* strains

Single knockout strains were generated using a transient CRISPR-Cas9 approach, with some modifications. All sgRNA expression cassettes were amplified from pV1093 plasmid in three PCR steps using sequentially the following three pairs of primers: SNR52/F and SNR52/R_GOI; sgRNA/F_GOI and sgRNA/R; and SNR52/N and sgRNA/N, where GOI represents the gene of interest. The Cas9 expression cassette was amplified from pV1093 plasmid with CaCas9/F and CaCas9/R primers. PCR reactions for both cassettes were performed according to Min *et al* (36) without any modifications. Repair templates were amplified from pV1093 plasmid using NAT_GOI_repair/F and NAT_GOI_repair/R primers. The PCR was performed in a total reaction volume of 50 μL: pV1093 plasmid (50 ng), 1 μL forward primer (10 μM), 1 μL reverse primer (10 μM), 25 μL CloneAMP mastermix (Takara), and distilled water up to 50 μL. Cycling conditions included an initial denaturation step at 98 °C for 2 min, followed by 40 cycles of denaturation at 98 °C for 30 s, annealing at 50 °C for 30 s, and extension at 72 °C for 1 min, and a final step at 72 °C for 10 min. *C. albicans* cells were transformed using the classical lithium acetate transformation method (37). The three cassettes were co-transformed in a single transformation (1 µg sgRNA cassette, 3 µg Cas9 cassette and 3 µg repair template). Positive transformants were selected in YPD supplemented with nourseothricin and confirmed by colony PCR with GOI-fwd and GOI- rv primers. Multiple-knockout strains and GFP fusions (38) were generated using the Hernday HIS-FLP system(39), with some modifications. For each strain, the specific gRNA expression cassette was obtained by cloning-free stitching PCR assembly of fragment A (amplified from pADH110 with AHO1096 and AH1098 primers) and fragment B (amplified from pADH147 with AHO1097 and Hernday-GOI primers) using AH01237 and AH01236 primers. The Cas9 cassette was obtained through digestion of pADH99 plasmid with MssI restriction enzyme. Both Cas9 and gRNA cassettes were co-transformed along with a linear DNA fragment (donor DNA) in a single transformation. The 220-bp deletion donors consisted of two 120-bp annealed oligonucleotides with homology to the upstream and downstream flanks of the targeted gene. Positive transformants were selected on YPD supplemented with nourseothricin and confirmed by colony PCR. After confirming the intended deletion, the NAT marker along with the Cas9 and gRNA expression cassettes were removed from the genome by inducing the expression of the maltose-inducible flipase (FLP) recombinase system on YP 2% maltose liquid medium overnight at 30 °C. Cells were then streaked on YPD agar for single-colony isolation and screened on YPD and YPD supplemented with nourseothricin to confirm efficient removal of the CRISPR components. All mutant strains were re-checked by colony PCR and validated by sequencing. Selected strains for *in vivo* assays were confirmed by whole-genome sequencing. All oligonucleotides used in this study are listed in Supplementary Table 2.

### Whole-genome sequencing

Genomic DNA samples were isolated using DNeasy UltraClean Microbial Kit (Qiagen), according to the manufacturer’s recommendations. Whole-genome sequencing was performed at Novogene (UK) using Illumina MiSeqII, with over ∼100× coverage. Raw sequencing reads were checked for quality using FastQC Software v.0.12.1 (https://www.bioinformatics.babraham.ac.uk/projects/fastqc/). Clean reads were assembled into scaffolds using MEGAHIT v.1.2.9 (40). Next, assembly quality was assessed using QUAST software v.5.2.0 (41). Gene prediction was performed with AUGUSTUS v3.5.0, using *C. albicans* SC5314 as the training model to improve accuracy. Genes of interest were aligned against each assembled genome using BLASTn with default parameters.

### Phenotypic assays in liquid and on solid media

Phenotypic assays were performed either on solid or liquid media. For evaluation of growth phenotypes by spot assays, cells were grown in solid YPD (2% w/v) medium. Cells were then harvested, washed twice in water and for each strain the optical density at 640 nm was adjusted to 1. A set of four 1:10 serial dilutions were performed and 3 µL of each suspension was spotted into the desired media, either SM or SC. Glucose (0.2% w/v) was used as a control carbon source. Cells were inoculated at 18 °C for 7 days, or at 37 °C for 2 days. For evaluation of growth rates, cells were pre-grown in SC media supplemented with 0.2% glucose (w/v) and then transferred to fresh media with an initial optical density of 0.01 at 640 nm. Optical densities were monitored every 2 hours at the same wavelength. The resulting data were processed using GraphPad Prism 9 Software. Data were log-transformed (base 10), and the exponential phase for each independent experiment was selected. The log of exponential growth equation was used for calculating growth rates.

### Radiolabelled Transport assays

*Candida* cells were directly grown in SC medium containing acetate (0.1% v/v, pH 6.0), collected at the optical density of 0.5 at 640 nm, washed twice in ice-cold deionized water, and resuspended in water to a final density of 20 – 40 mg (dry weight)/mL. For each reaction, a cell suspension of 30 µL was mixed with 60 µL of potassium phosphate buffer (100 mM, pH 6.0). After 2 min of incubation at 30 °C, each reaction was started by the addition of 10 µL of radiolabelled [1-^14^C]-acetic acid (sodium salt, 55.2 mCi/mmol, Perkin Elmer) at the desired concentration for 15 s. Reactions were stopped by dilution with 100 µL ice-cold acetic acid (100 mM, pH 6.0). The reaction mixtures were kept on ice, centrifuged at 13 000 rpm, washed with 1 mL ice-cold deionized water and the final pellet was resuspended in 1 mL of scintillation fluid (Opti-phase HiSafe II). Radioactivity was measured in a Packard Tri-Carb 2200 CA liquid scintillation counter. Each reaction was prepared in duplicate, and each assay was repeated three times. The transport kinetics best fitting the experimental initial uptake rates and the kinetic parameters were determined by a computer-assisted nonlinear regression analysis (GraphPad Software, Prism 9).

### Fluorescence Microscopy

*C. albicans* cells were grown at 30 °C in SM medium containing glucose (0.2% w/v) until mid-exponential phase (OD_640 nm_ = 0.5), washed twice with deionized water and then transferred to fresh SC medium supplemented with acetate (0.1% v/v, pH 6.0) for 6, 8 and 24 h. Cells were examined with a Leica Microsystems DM-5000B epifluorescence microscope with appropriate filter settings. Images were acquired with a Leica DCF350FX digital camera and processed with LAS AF Leica Microsystems software. Pictures are representative of three independent experiments.

### Murine model of gastrointestinal colonisation

All animal procedures were performed in accordance with institutional guidelines of KU Leuven. The specific protocols for this study were approved by the KU Leuven Ethical Committee (project P010/2020). The laboratory animal usage license number is LA1210204. Female C57BL6 mice (aged 8-12 weeks) were obtained from Charles River laboratories. No statistical methods were used to predetermine group size. Each group of mice (n = 4) was assigned at random, housed separately in individual ventilated autoclave-sterilized cages, provided with food and water *ad libitum* and adapted to standardized environmental conditions (temperature, 23 ± 2 °C; humidity, 55 ± 10%; 12 h light/dark cycles) for at least 1 week before the beginning of each experiment. Four groups of mice were provided with sterile drinking water containing 2 mg/mL streptomycin and 1 mg/mL ampicillin four days before exposure to *C. albicans* strains and maintained on antibiotic treatment until the end of the experiment. As a control experiment, another four groups of mice did not receive any treatment. Each experiment (except control, untreated groups) was repeated twice (n = 8). For experimental colonisation following oral gavage, *C. albicans* strains were grown overnight in YPD medium and washed twice in sterile phosphate-buffered saline (PBS). Cell densities were adjusted with PBS and confirmed by flow cytometry-based cell counting and by plating (CFUs). Mice were gavaged with 5×10^7^ *C. albicans* cells and sacrificed after seven days of exposure. Faecal samples were retrieved daily before and after *C. albicans* colonisation to assay fungal burdens (CFUs) and microbiome composition (16S rRNA gene amplicon sequencing). Gut compartments (small intestine, cecum and colon) from each mouse were harvested in PBS, weighted, homogenized with glass beads using a FastPrep instrument (MP Biomedicals) and then split into samples to determine fungal burdens (CFUs) and SCFA concentrations. For CFU counting, both faecal and tissue samples were serially diluted tenfold, 50 µL of each dilution plated on CHROMagar, and plates grown overnight at 37 °C. Fungal burdens were expressed as CFU per gram of feces or tissue, respectively.

### 16S rRNA gene amplicon sequencing

Total genomic DNA was isolated from stool samples using QIAamp PowerFecal Pro DNA Kit (Quiagen) according to manufacturer’s instructions. The DNA concentration and purity were determined with NanoDrop 2000 UV-Vis spectrophotometer (Thermo Scientific). The hypervariable region V3-V4 of the bacterial 16S rRNA gene was amplified with primer pairs 338F: 5’-ACTCCTACGGGAGGCAGCA-3’ and 806R: 5’-GGACTACHVGGGTWTCTAAT-3’. Both the forward and reverse 16S primers were tailed with sample-specific Illumina index sequences to allow for deep sequencing. The PCR was performed in a total reaction volume of 10 μL: 5-50 ng DNA template, 0.3 μL forward primer (10 μM), 0.3 μL reverse primer (10 μM), 5 μL KOD FX Neo Buffer, 2 μL dNTP (2 mM each), 0.2 μL KOD FX Neo, and finally ddH_2_O up to 20 μL. Cycling conditions included an initial denaturation step at 95 °C for 5 min, followed by 20 cycles of denaturation at 95 °C for 30 s, annealing at 50 °C for 30 s, and extension at 72 °C for 40 s, and a final step at 72 °C for 7 min. The amplified products were purified with Omega DNA purification kit (Omega) and quantified using Qsep-400 (BiOptic). The amplicon library was paired-end sequenced (2×250) on an Illumina novaseq6000 (Beijing Biomarker Technologies Co). The bioinformatic analysis was performed with the aid of the BMKCloud (http://www.biocloud.net/). According to quality of single nucleotide, raw data was primarily filtered by Trimmomatic v.033 (42). Identification and removal of primer sequences was process by Cutadapt v.1.9.1 (43). PE reads obtained from previous steps were assembled by USEARCH (44) and followed by chimera removal using UCHIME (45). The high-quality reads generated from above steps were used in the following analysis. Sequences with similarity >97% were clustered into the same operational taxonomic unit (OTU) by USEARCH (44), and the OTUs counts less than 2 in all samples were filtered. Clean reads then were conducted on feature classification to output an ASVs (amplicon sequence variants) by dada2 (46), and the ASVs counts less than 2 in all samples were filtered. Taxonomy annotation of the OTUs was performed based on the Naive Bayes classifier in QIIME2 (47) using the SILVA database (48) with a confidence threshold of 70%. Alpha diversity was calculated and displayed by the QIIME2 and R software, respectively. Beta diversity was determined to evaluate the degree of similarity of microbial communities from different samples using QIIME. Principal coordinate analysis (PCoA), heatmaps, UPGMA and nonmetric multidimensional scaling (NMDS) were used to analyse the beta diversity. Furthermore, we employed Linear Discriminant Analysis (LDA) effect size (LEfSe) (49) to test the significant taxonomic difference among groups. A logarithmic LDA score of 4.0 was set as the threshold for discriminative features. To explore the dissimilarities of the microbiome among different factors, a redundancy analysis (RDA) was performed in R using the package ‘vegan’.

### SCFAs extraction and quantification

SCFAs acids were extracted from gut homogenates colonised by WT *C. albicans* (small intestine, cecum and colon). After homogenization with glass beads using a FastPrep instrument (MP Biomedicals), samples were centrifuged at 15000 rpm for 10 min and the supernatants taken for analysis. For each sample, 2-ethyl butyrate was used as an internal standard at a final concentration of 5 mM. A mixture of SCFAs, 10 mM each (lactate, acetate, propionate, formate and butyrate), was used as an external standard. SCFAs were then extracted by the addition of 0.1 mL HCl (37%) and 0.4 mL diethyl ether to 0.2 mL of each supernatant followed by 1 min of vortex mixing. After centrifugation at 3000 g for 10 min, the ether layer was removed and transferred to a separate capped vial. A further 0.4 mL diethyl ether was added to the aqueous layer and a second extraction performed. The ether extracts were combined, and 40 µL *N*-methyl-*N*-t-butyldimethylsilyltrifluoroacetamide (MTBSTFA) added before heating at 80 °C for 20 min. SCFAs were quantified simultaneously as their tertiary butyldimethylsilyl (t-BDMS) derivatives using a gas chromatography-mass spectrometer (Trace 1300 – ISQ QD equipped with a TriPlus RSH autosampler and a Restek Rxi-5ms capillary GC column 30 m x 0.25 mmID). Helium was used as carrier gas with a flow rate of 1.4 mL/min. Injection was carried out at 250 °C in split mode after 1 min and with a ratio of 1:10. The temperature was first held at 50 °C for 1 min and then allowed to rise to 260 °C at a rate of 50 °C/min, followed by a second ramp of 2 °C/min until 325 °C was reached; that temperature was maintained for 3 min. The mass detector was operated in scan mode (50 to 600 atomic mass units), using electron impact ionization (70 eV). The temperatures of the MS transfer line and detector were 325 °C and 250 °C, respectively. SCFAs were identified by their retention time relative to the internal standard and specific mass spectrometric patterns. Peak areas obtained from the analysis of the external standard solution, to which internal standard had been added, were used to calculate the relative response factors for each acid with respect to the internal standard. Final concentrations were determined relative to both internal and external standards, and then normalized against tissue weight.

### Statistics and reproducibility

All repeated independent experiments showed similar results. No statistical method was used to predetermine sample sizes, and no data were excluded from the analyses. Statistical analyses were performed using GraphPad Prism 9 software. All data are presented as mean ± sd unless otherwise stated. All details of the statistics, including the specific tests used for each experiment, are provided in the corresponding figure legends. All schematic representations and figures were created or assembled using BioRender (https://biorender.com).

## Acknowledgments

This work was supported by the MetaFungal project PTDC/BIA-MIC/5246/2020 (https://doi.org/10.54499/PTDC/BIA-MIC/5246/2020), funded by the Portuguese Foundation for Science and Technology. Work at CBMA was supported by Contrato Programa UIDB/04050/2020 (https://doi.org/10.54499/UIDB/04050/2020). RA acknowledges FEBS for a short-term fellowship at University of Exeter (UK) and FEMS for her stay at KU Leuven (Belgium) in the scope of the FEMS-Jensen Award. FG, PA and AG acknowledge FCT for their PhD fellowships (2023.03135.BD; 2024.03178.BDANA, 2021.08564.BD, respectively). Work at KU Leuven was supported by the Research Council (grant #C14/22/075) and the Fund for Scientific Research Flanders (FWO grant #G0C0622N). AJPB was supported by a programme grant from the UK Medical Research Council (MR/M026663/2), a Wellcome Investigator Award (224323/Z/21/Z), and by the Medical Research Council Centre for Medical Mycology at the University of Exeter (MR/N006364/2). The funders had no role in study design, data collection and analysis, decision to publish, or preparation of the manuscript. For the purpose of open access, the author has applied a CC BY public copyright licence to any Author Accepted Manuscript version arising from this submission.

## Author Contributions

RA performed the experiments and was responsible for the formal analysis, curation, and visualisation of the data. FG, CBA, VF, PA, AG, and QM assisted with the in vitro experiments, while WVG, SW, OVG, and RV assisted with the in vivo experiments. RD contributed to the whole genome sequence analysis. ISS and MC contributed to the transport data analysis. SP, PVD, and AJPB contributed to funding acquisition and were major contributors in conceptualization, validation and supervision. RA drafted the original manuscript with additional input from all authors.

