## Supplemental data for "The *ATO* gene family governs *Candida albicans* colonisation in the dysbiotic gastrointestinal tract"

Supplementary material includes Figures S1 and S2 and Tables S1 and S2.

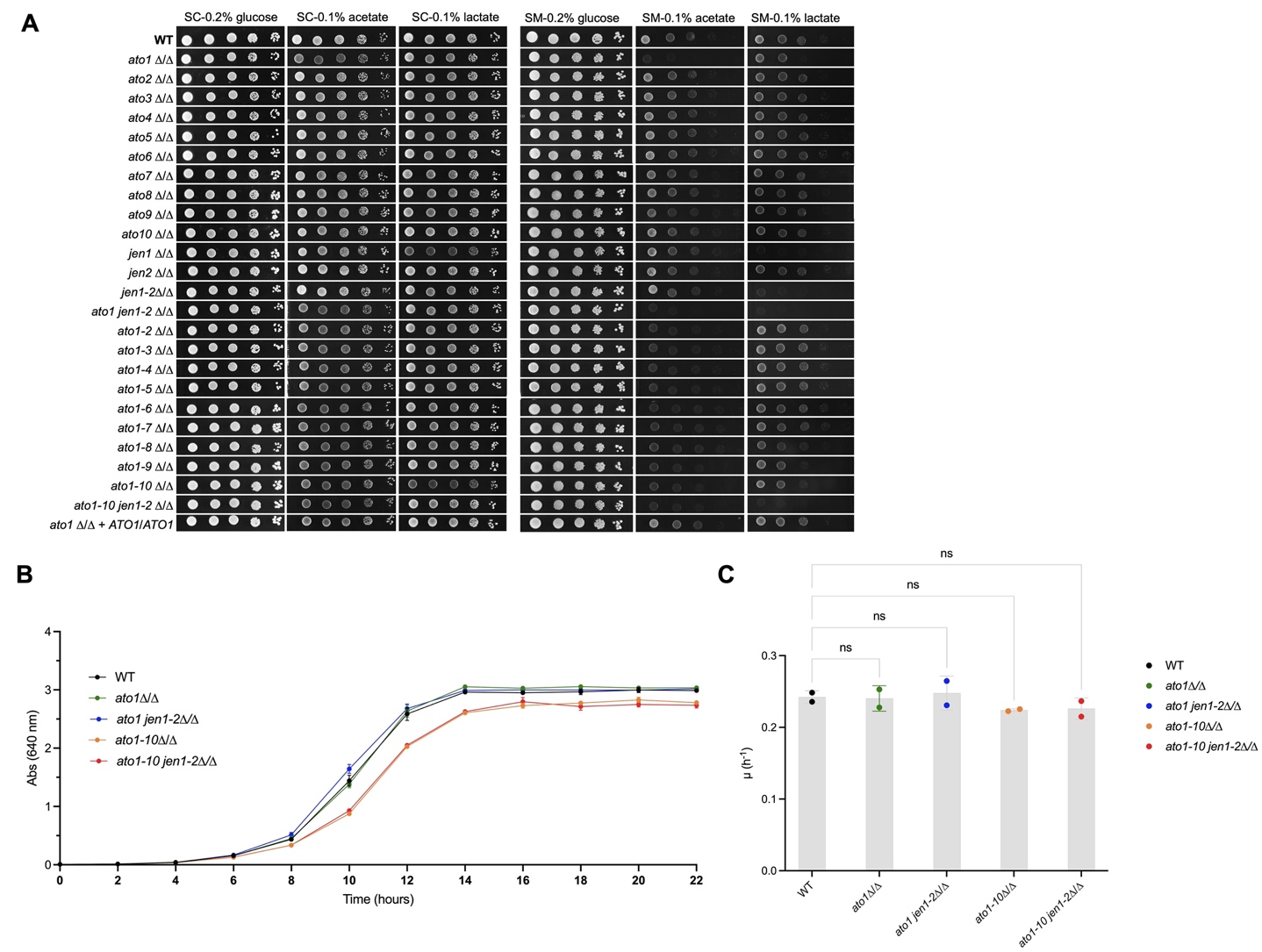

**Figure S1. *Candida albicans* growth phenotypes. Related to Figure 2.**

**A.** Growth phenotypes of *C. albicans* WT and knockout strains in Synthetic Complete (SC) and Synthetic Minimal (SM) media supplemented with specific defined carbon sources: glucose (0.2% w/v), acetate (0.1% v/v, pH 6.0), lactate (0.1% v/v, pH 5.0). Cells were serially diluted, spotted on solid media and incubated at 37 ºC for 2 days. **B**. Growth assays of *C. albicans* strains used in the gut colonisation experiments. Cells were grown in SC liquid medium supplemented with glucose 0.2% (w/v) and incubated at 30 °C for 24 hours. Data are shown as mean ± SEM of two independent biological experiments repeated in duplicate. “ns” indicates non-significance between strains, as determined by one-way ANOVA using Tukey’s multiple comparisons test. **C**. Statistical analyses of the differences in growth rates (h-1) of *C. albicans* strains observed in B. “ns” indicates non-significant.

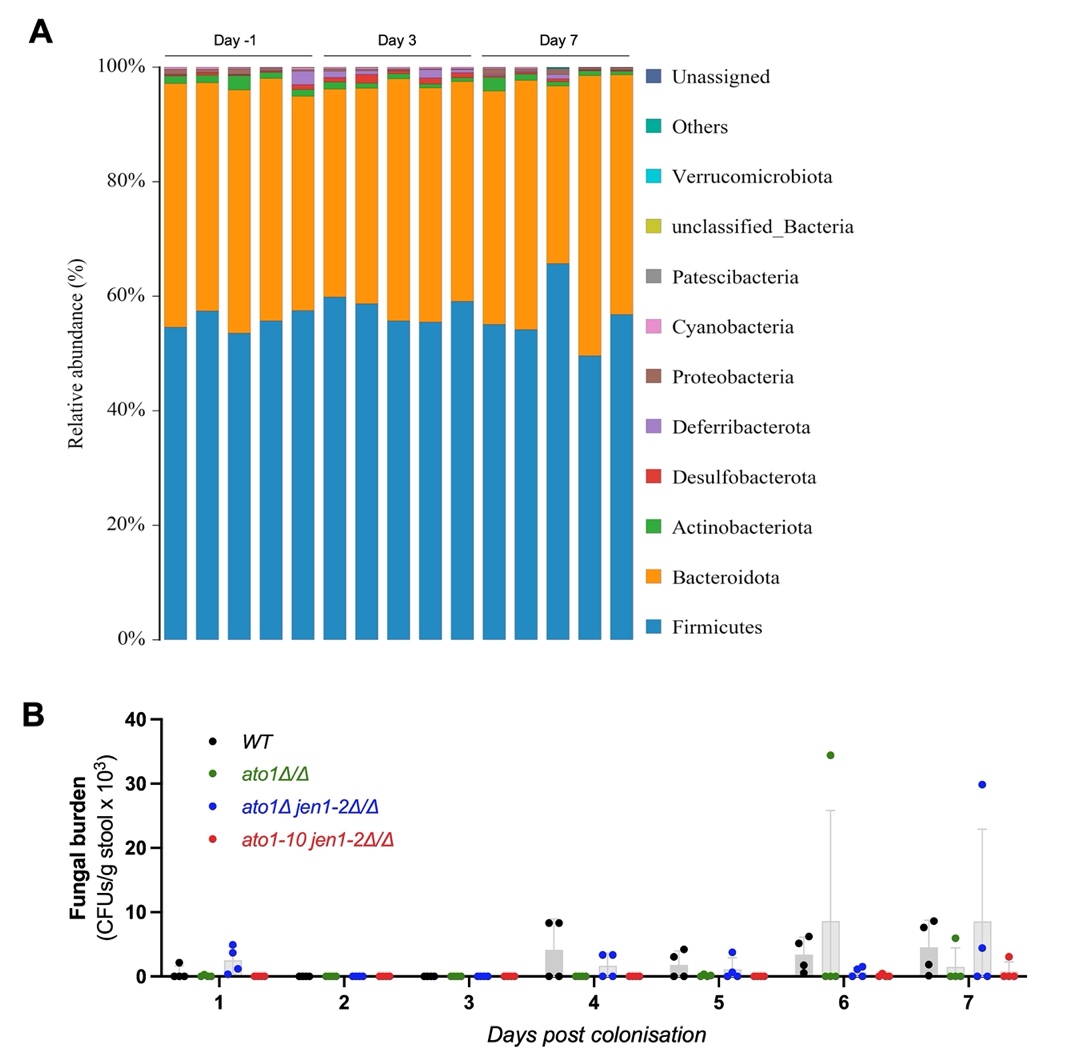

**Figure S2. *Candida albicans* colonisation in the gut.** **Related to Figure 3.**

**A.** Relative abundance of taxa (%) in untreated mice before (Day -1) and after *C. albicans* colonisation (Day 3 and 7). Each bar represents one independent group of 4 mice. **B.** Fecal colonisation levels (CFUs/g stool) of *C. albicans* strains (*n* = 4 per strain) in untreated mice over time.

**Table S1**. List of strains used in this study.

| **Strain**^a^ | **Genotype** | **Reference** |
| --- | --- | --- |
| WT | SC5314 | N/A |
| *ato1*∆/∆ | *ATO1*::NAT/*ATO1*::NAT | This study |
| *ato2*∆/∆ | *ATO2*::NAT/*ATO2*::NAT | This study |
| *ato3*∆/∆ | *ATO3*::NAT/*ATO3*::NAT | This study |
| *ato4*∆/∆ | *ATO4*::NAT/*ATO4*::NAT | This study |
| *ato5*∆/∆ | *ATO5*::NAT/*ATO5*::NAT | This study |
| *ato6*∆/∆ | *ATO6*::NAT/*ATO6*::NAT | This study |
| *ato7*∆/∆ | *ATO7*::NAT/*ATO7*::NAT | This study |
| *ato8*∆/∆ | *ATO8*::NAT/*ATO8*::NAT | This study |
| *ato9*∆/∆ | *ATO9*::NAT/ATO9::NAT | This study |
| *ato10*∆/∆ | *ATO10*::NAT/*ATO10*::NAT | This study |
| *jen1*∆/∆ | *JEN1*::NAT/*JEN1*::NAT | This study |
| *jen2*∆/∆ | *JEN2*::NAT/*JEN2*::NAT | This study |
| *ato1*∆/∆ | *ato1*∆/∆ | This study |
| *ato1-2*∆/∆ | *ato1*∆/∆ *ato2*∆/∆ | This study |
| *ato1-3*∆/∆ | *ato1*∆/∆ *ato2*∆/∆ *ato3*∆/∆ | This study |
| *ato1-4*∆/∆ | *ato1*∆/∆ *ato2*∆/∆ *ato3*∆/∆ *ato4*∆/∆ | This study |
| *ato1-5*∆/∆ | *ato1*∆/∆ *ato2*∆/∆ *ato3*∆/∆ *ato4*∆/∆ *ato5*∆/∆ | This study |
| *ato1-6*∆/∆ | *ato1*∆/∆ *ato2*∆/∆ *ato3*∆/∆ *ato4*∆/∆ *ato5*∆/∆ *ato6*∆/∆ | This study |
| *ato1-7*∆/∆ | *ato1*∆/∆ *ato2*∆/∆ *ato3*∆/∆ *ato4*∆/∆ *ato5*∆/∆ *ato6*∆/∆ *ato7*∆/∆ | This study |
| *ato1-8*∆/∆ | *ato1*∆/∆ *ato2*∆/∆ *ato3*∆/∆ *ato4*∆/∆ *ato5*∆/∆ *ato6*∆/∆ *ato7*∆/∆ *ato8*∆/∆ | This study |
| *ato1-9*∆/∆ | *ato1*∆/∆ *ato2*∆/∆ *ato3*∆/∆ *ato4*∆/∆ *ato5*∆/∆ *ato6*∆/∆ *ato7*∆/∆ *ato8*∆/∆ *ato9*∆/∆ | This study |
| *ato1-10*∆/∆ | *ato1*∆/∆ *ato2*∆/∆ *ato3*∆/∆ *ato4*∆/∆ *ato5*∆/∆ *ato6*∆/∆ *ato7*∆/∆ *ato8*∆/∆ *ato9*∆/∆ *ato10*∆/∆ | This study |
| *jen1-2*∆/∆ | *jen1*∆/∆ *jen2*∆/∆ | This study |
| *ato1 jen1*∆/∆ | *ato1*∆/∆ *jen1*∆/∆ | This study |
| *ato1* *jen1-2*∆/∆ | *ato1*∆/∆ *jen1*∆/∆ *jen2*∆/∆ | This study |
| *ato1-10 jen1-2*∆/∆ | *ato1-10*∆/∆ *jen1*∆/∆ *jen2*∆/∆ | This study |
| *ATO1*-GFP | *ATO1*-GFP/*ATO1*-GFP | This study |

^a^ For constructed mutants, three independent transformants were considered individual strains (biological repeats).

**Table S2**. List of oligonucleotides used for strain engineering and screening.

| **Name** | **Sequence** |
| --- | --- |
| SNR52/F | AAGAAAGAAAGAAAACCAGGAGTGAA |
| SNR52/R_ATO1 | ccaaatgcttcaactaaatcCAAATTAAAAATAGTTTACGCAAGTC |
| SNR52/R_ATO2 | ccgaaagcagccatcaattcCAAATTAAAAATAGTTTACGCAAGTC |
| SNR52/R_ATO3 | ccgaaggcagccatcaagtcCAAATTAAAAATAGTTTACGCAAGTC |
| SNR52/R_ATO4 | tgctgctattggtttagtttCAAATTAAAAATAGTTTACGCAAGTC |
| SNR52/R_ATO5 | tcaccagttatcgtacacgtCAAATTAAAAATAGTTTACGCAAGTC |
| SNR52/R_ATO6 | tggtgtttgttttgttatttCAAATTAAAAATAGTTTACGCAAGTC |
| SNR52/R_ATO7 | tcaccagcataagagacagtCAAATTAAAAATAGTTTACGCAAGTC |
| SNR52/R_ATO8 | tcaccagaaatctcaactttCAAATTAAAAATAGTTTACGCAAGTC |
| SNR52/R_ATO9 | gcagcagtgaccaatattccCAAATTAAAAATAGTTTACGCAAGTC |
| SNR52/R_ATO10 | tcaccagtagttctaactctCAAATTAAAAATAGTTTACGCAAGTC |
| SNR52/R_JEN1 | caagtaacatcagtaacactCAAATTAAAAATAGTTTACGCAAGTC |
| SNR52/R_JEN2 | ctggcctttttaggagcatcCAAATTAAAAATAGTTTACGCAAGTC |
| sgRNA/F_ATO1 | gatttagttgaagcatttggGTTTTAGAGCTAGAAATAGCAAGTTAAA |
| sgRNA/F_ATO2 | gaattgatggctgctttcggGTTTTAGAGCTAGAAATAGCAAGTTAAA |
| sgRNA/F_ATO3 | gacttgatggctgccttcgggTTTTAGAGCTAGAAATAGCAAGTTAAA |
| sgRNA/F_ATO4 | aaactaaaccaatagcagcaGTTTTAGAGCTAGAAATAGCAAGTTAAA |
| sgRNA/F_ATO5 | acgtgtacgataactggtgaGTTTTAGAGCTAGAAATAGCAAGTTAAA |
| sgRNA/F_ATO6 | aaataacaaaacaaacaccaGTTTTAGAGCTAGAAATAGCAAGTTAAA |
| sgRNA/F_ATO7 | actgtctcttatgctggtgaGTTTTAGAGCTAGAAATAGCAAGTTAAA |
| sgRNA/F_ATO8 | aaagttgagatttctggtgaGTTTTAGAGCTAGAAATAGCAAGTTAAA |
| sgRNA/F_ATO9 | ggaatattggtcactgctgcGTTTTAGAGCTAGAAATAGCAAGTTAAA |
| sgRNA/F_ATO10 | agagttagaactactggtgaGTTTTAGAGCTAGAAATAGCAAGTTAAA |
| sgRNA/F_JEN1 | agtgttactgatgttacttgGTTTTAGAGCTAGAAATAGCAAGTTAAA |
| sgRNA/F_JEN2 | gatgctcctaaaaaggccagGTTTTAGAGCTAGAAATAGCAAGTTAAA |
| sgRNA/R | ACAAATATTTAAACTCGGGACCTGG |
| SNR52/N | GCGGCCGCAAGTGATTAGACT |
| sgRNA/N | GCAGCTCAGTGATTAAGAGTAAAGATGG |
| CaCas9/F | ATCTCATTAGATTTGGAACTTGTGGGTT |
| CaCas9/R | TTCGAGCGTCCCAAAACCTTCT |
| NAT_ATO1_repair/F | ACACTACAATAAACTTTTAACAACACACATAAATAACTAACTACTACAACTACAACTACAACCACTACACTTATCAAATCagtctaatcacttgcggccgc |
| NAT_ATO1_repair/R | TCATCATTAAAAAAAAAAATCAATACCAGTATTGATTGATTGATTGATTGATTGATTGATTGATTACTTGGCCAATTATTggaccacctttgattgtaaatag |
| NAT_ATO2_repair/F | CCAAATTGCTGTTCATTTGTTTTAGCTATTATTTTGTTCGTTCTACGAAGAATCTTATTATTGATCTACTGTTAAAGAATagtctaatcacttgcggccgc |
| NAT_ATO2_repair/R | TTATAGTTGAATCGTACAATTGAACAATCAAATTAAGCATTAAAATTAGATAAGTAGCAACAACTGAAAGAGTTTTGCCAggaccacctttgattgtaaatag |
| NAT_ATO3_repair/F | TTCGAAATAGAAAAAAACTGTTTCTTTTATATAAAGTTCTACTATCTATATCCAATATCGATAAATAAAAGGAAACAACTagtctaatcacttgcggccgc |
| NAT_ATO3_repair/R | AATTATAGTAGCAATAAAGAAAAAAAAAATCAAATGATAGCCATGGTGAATGACAATAAAATGCAGCTAGACAGATATTTggaccacctttgattgtaaatag |
| NAT_ATO4_repair/F | ACACGGCTATAAAAGGCCAACCATCTATCCATATAGATACCAGCTTTAATTACCACTACAAATCCAACAAAATTAATCAAagtctaatcacttgcggccgc |
| NAT_ATO4_repair/R | ACCGCAACTGCAACTTGAAAGTACAGAAAGTACACAATACTGAATCTTTAGGGTTATTGTAGCCGGGTTCCGTTTTTGAGggaccacctttgattgtaaatag |
| NAT_ATO5_repair/F | CAATCTACCGATAATAAAAGACCCTATTTCCCTATTACCCAAATTTTATTAAAAGCCAAGTTAACAAAACATACAAAACAagtctaatcacttgcggccgc |
| NAT_ATO5_repair/R | ATAAATTAATATTTAATACATTTGTGGCTTATAAAAAATGTTTCCAGAACAAGTTACAAACAAAAATATAACGTTATACTggaccacctttgattgtaaatag |
| NAT_ATO6_repair/F | TAGTTAAGTTTAGGTGTTCTCATTTTAATCAACATATACACATATTAGAAACCCATAATCACAACTACACTATCACAATCagtctaatcacttgcggccgc |
| NAT_ATO6_repair/R | AAGGTAACAAAAGAGAAAGAAAGAATATTAACAGACATGAATAATAGAATTCTAGGATGCACAGCATAGCTAGTTTAGATggaccacctttgattgtaaatag |
| NAT_ATO7_repair/F | ATAAAGCTATAAATTGCCTGTGTTTTCACCTTTCATAGCTATCATACGTTCTATACTCAGACCATCCATCCTCCTATAACagtctaatcacttgcggccgc |
| NAT_ATO7_repair/R | TACTTGTTCTAATAAATAATAAATCAATTTATACCAGACAAGTTGACCTCCCCCTTTTTCTGTAGAACTCAGATCTCATCggaccacctttgattgtaaatag |
| NAT_ATO8_repair/F | TTGGTAATCAGTATTTTTTGAATGTTCCATTTGCAAGAGAAATACTTGACCAAAAAAGTTATAAAATCAAATTATAAGTAagtctaatcacttgcggccgc |
| NAT_ATO8_repair/R | TAAATATTAAAGAAATTATTAAAAAAAAATAGAAAAATGAAAGAAATCCAAATAAACCGTTTTCCAGCTCTTAACCTTTGggaccacctttgattgtaaatag |
| NAT_ATO9_repair/F | TTAAAATATATAACTATATAAAATATTTGACATTTCTTATTCCTTTTATACCAAACTCATGTTTCACGTCCTTCCGGCATagtctaatcacttgcggccgc |
| NAT_ATO9_repair/R | TTGTTATTGTTGTCCCTTAAAAAATGATAAAAAAAAAGTTTGTTGTCTTCCCCCTTTGATTGTAGTTTTCCACTAACTTCggacctttgattgtaaatag |
| NAT_ATO10_repair/F | AATTGATAGTTTATTCTTTTTTGTAATCAATAGTTTTGTTCAAACATTAAGCAACACAATATACTATACCAAAATTTCTTagtctaatcacttgcggccgc |
| NAT_ATO10_repair/R | AGAACGGGCTATATCATCATTTAAGTCAGACATTACTAATTTACGCTAAGGTGTTAGCTTATTGGACCAACTACTTTAAAggacctttgattgtaaatag |
| NAT_JEN1_repair/F | CGTATTTTCCTTCTTAATATTCTTCAATACTTTTTATAAATATTAAGATAAACTCCATATCACACCTACACACACACAATagtctaatcacttgcggccgc |
| NAT_JEN1_repair/R | CCTAAATATAAATAATCCGTTTATTCATGTCATCATTGAATTAGAGCTTATTGTTTATATATTTATATAATTTTATGGTAggacctttgattgtaaatag |
| NAT_JEN2_repair/F | GTTATAAAAACCCAATACATCACATTACATTATAAATCATACACATATACATATATATAATAACTAACCATAGAAATATTagtctaatcacttgcggccgc |
| NAT_JEN2_repair/R | CATAAACCCATTTATTATCAAAATAAACTATACTTGTTATATGATAATCTAAACATAAATAAATAAAATACCACCAACACggacctttgattgtaaatag |
| ATO1-fwd | CAACTACAACTACAACCACT |
| ATO1-rv | CTCGGAGTTTTTCAATACGT |
| ATO2-fwd | CGTCGTCTCCCTTCTATTCT |
| ATO2-rv | TGTCTAGTTGGTTCCCGTCC |
| ATO3-fwd | CTTCTCTCGTTCCGTTTTCT |
| ATO3-rv | CAAATGATAGCCATGGTGAA |
| ATO4-fwd | GGCTATAAAAGGCCAACCAT |
| ATO4-rv | TCGTCGTCAGGTAATTGCAG |
| ATO5-fwd | GGATGAACAGCCTTATTGTT |
| ATO5-rv | GTTCGATCATCAACCTTCAC |
| ATO6-fwd | TTGTAGTTTCACTCAGGTTT |
| ATO6-rv | TCCATGAGTTACCTCCACTG |
| ATO7-fwd | GTGTTTTCACCTTTCATAGC |
| ATO7-rv | ATTGAGTGTAGTTGTCTCTG |
| ATO8-fwd | GTTACAATTGCAAACTGCTT |
| ATO8-rv | AAGTAGTCGTGCATGTTTTC |
| ATO9-fwd | CGTTCTAAATCTGTGTAGGC |
| ATO9-rv | GCTAATTATGAGATAGCTTC |
| ATO10-fwd | CCAATACCACTCTTTAATGT |
| ATO10-rv | GGAAGGACGTGAAACATGAG |
| JEN1-fwd | TTATGCCTGTTTACCAATCC |
| JEN1-rv | CTACCATAAATACACGACTG |
| JEN2-fwd | CCCAATATCTTTGGTTTGCT |
| JEN2-rv | TAAGATGGCGTGGTAGCTTA |
| AHO1096 | GACGGCACGGCCACGCGTTTAAACCGCC |
| AHO1098 | caaattaaaaatagtttacgcaag |
| AHO1097 | CCCGCCAGGCGCTGGGGTTTAAACACCG |
| Hernday-ATO1 | CGTAAACTATTTTTAATTTGgatttagttgaagcatttggGTTTTAGAGCTAGAAATAG |
| Hernday-ATO2/3 | CGTAAACTATTTTTAATTTGacatcttctcaaaaatctgtGTTTTAGAGCTAGAAATAG |
| Hernday-ATO4/5 | CGTAAACTATTTTTAATTTGgaattgatgcaagcatttggGTTTTAGAGCTAGAAATAG |
| Hernday-ATO6/7 | CGTAAACTATTTTTAATTTGatttggtaatgcttcagcactGTTTTAGAGCTAGAAATAG |
| Hernday-ATO8 | CGTAAACTATTTTTAATTTGaaagttgagatttctggtgaGTTTTAGAGCTAGAAATAG |
| Hernday-ATO9/10 | CGTAAACTATTTTTAATTTGacatcactcaaagatatcgaGTTTTAGAGCTAGAAATAG |
| Hernday-JEN1 | CGTAAACTATTTTTAATTTGGCAAACAATACTAATCACAAGTTTTAGAGCTAGAAATAG |
| Hernday-JEN2 | CGTAAACTATTTTTAATTTGTGCTCCTAAAAAGGCCAGAGGTTTTAGAGCTAGAAATAG |
| AHO1237 | aggtgatgctgaagctattgaag |
| AHO1236 | TAAAGCTGCCACAAGAGGTATTTC |
| Delete-ATO1-fw | ATCAATATCAATATCAATAGACACTACAATAAACTTTTAACAACACACATAAATAACTAACTACTACAACTACAACTACAACCACTACACTTATCAAATCAATAATTGGCCAAGTAATCA |
| Delete-ATO1-rv | TTTGGGAACCATTAAATAAATAAATAAAAAAATCAAATCTTCATCATTAAAAAAAAAAATCAATACCAGTATTGATTGATTGATTGATTGATTGATTGATTGATTACTTGGCCAATTATT |
| Delete-ATO2-fw | TCTTTGTATATCCAAGTTGGCCAAATTGCTGTTCATTTGTTTTAGCTATTATTTTGTTCGTTCTACGAAGAATCTTATTATTGATCTACTGTTAAAGAATTGGCAAAACTCTTTCAGTTG |
| Delete-ATO2-rv | AGTGGTTCCCGTCCCCCAAGATATATATAGATAAAATTTATAGTTGAATCGTACAATTGAACAATCAAATTAAGCATTAAAATTAGATAAGTAGCAACAACTGAAAGAGTTTTGCCA |
| Delete-ATO3-fw | CTATAAGTGTGTTCTTTTCTTTCGAAATAGAAAAAAACTGTTTCTTTTATATAAAGTTCTACTATCTATATCCAATATCGATAAATAAAAGGAAACAACTAAATATCTGTCTAGCTGCAT |
| Delete-ATO3-rv | CCCAGTCCCAATATAAAACAACACAATAAATAAATAAATAAATTATAGTAGCAATAAAGAAAAAAAAAATCAAATGATAGCCATGGTGAATGACAATAAAATGCAGCTAGACAGATATTT |
| Delete-ATO4-fw | TTTCATCCCTTGAAGATAATACACGGCTATAAAAGGCCAACCATCTATCCATATAGATACCAGCTTTAATTACCACTACAAATCCAACAAAATTAATCAACTCAAAAACGGAACCCGGCT |
| Delete-ATO4-rv | AAACACACAAAGATAAATCTTACAAATTGCAACGAGGATGACCGCAACTGCAACTTGAAAGTACAGAAAGTACACAATACTGAATCTTTAGGGTTATTGTAGCCGGGTTCCGTTTTTGAG |
| Delete-ATO5-fw | CGGGGTGTATAAATACCCAGCAATCTACCGATAATAAAAGACCCTATTTCCCTATTACCCAAATTTTATTAAAAGCCAAGTTAACAAAACATACAAAACAAGTATAACGTTATATTTTTG |
| Delete-ATO5-rv | TCATTTCAATGAAATAAGGGGGGGAGAGGGGGGGGGGGCGATAAATTAATATTTAATACATTTGTGGCTTATAAAAAATGTTTCCAGAACAAGTTACAAACAAAAATATAACGTTATACT |
| Delete-ATO6-fw | TTAATCCATACATACTACTTTAGTTAAGTTTAGGTGTTCTCATTTTAATCAACATATACACATATTAGAAACCCATAATCACAACTACACTATCACAATCATCTAAACTAGCTATGCTGT |
| Delete-ATO6-rv | TTATTGCTAAAAAAAAAAAAAAGTACAAATCAATTATACGAAGGTAACAAAAGAGAAAGAAAGAATATTAACAGACATGAATAATAGAATTCTAGGATGCACAGCATAGCTAGTTTAGAT |
| Delete-ATO7-fw | GTCTTTATAAAAATCCACATATAAAGCTATAAATTGCCTGTGTTTTCACCTTTCATAGCTATCATACGTTCTATACTCAGACCATCCATCCTCCTATAACGATGAGATCTGAGTTCTACA |
| Delete-ATO7-rv | ATAAAAACCTAAACAAATTGTACATTTTGTGGACTTATTGTACTTGTTCTAATAAATAATAAATCAATTTATACCAGACAAGTTGACCTCCCCCTTTTTCTGTAGAACTCAGATCTCATC |
| Delete-ATO8-fw | CAGCTCAGTTATAAATAGGTTTGGTAATCAGTATTTTTTGAATGTTCCATTTGCAAGAGAAATACTTGACCAAAAAAGTTATAAAATCAAATTATAAGTACAAAGGTTAAGAGCTGGAAA |
| Delete-ATO8-rv | AAATTATGTATTAAGTATAGAACACAATAATTAATGATTATAAATATTAAAGAAATTATTAAAAAAAAATAGAAAAATGAAAGAAATCCAAATAAACCGTTTTCCAGCTCTTAACCTTTG |
| Delete-ATO9-fw | AAACATTTAATTTCCCATAATTAAAATATATAACTATATAAAATATTTGACATTTCTTATTCCTTTTATACCAAACTCATGTTTCACGTCCTTCCGGCATGAAGTTAGTGGAAAACTACA |
| Delete-ATO10-rv | ACCATTGCCTACACAGATTTAGAACGGGCTATATCATCATTTAAGTCAGACATTACTAATTTACGCTAAGGTGTTAGCTTATTGGACCAACTACTTTAAATGTAGTTTTCCACTAACTTC |
| Delete-ATO9/10-fw | AACAATAGAAATGCCGCTGGAATTGATAGTTTATTCTTTTTTGTAATCAATAGTTTTGTTCAAACATTAAGCAACACAATATACTATACCAAAATTTCTTGAAGTTAGTGGAAAACTACA |
| Delete-ATO9/10-rv | TAGAAATATAGAAGGCTCCCGTCTCCCACCCCATAATGTGTTGTTATTGTTGTCCCTTAAAAAATGATAAAAAAAAAGTTTGTTGTCTTCCCCCTTTGATTGTAGTTTTCCACTAACTTC |
| Delete-JEN1-fw | ATAAATATTAAGATAAACTCCATATCACACCTACACACACACAATTACCATAAAATTATATAAATATATAAACAATAAGCTCTAATTCA |
| Delete-JEN1-rv | TGAATTAGAGCTTATTGTTTATATATTTATATAATTTTATGGTAATTGTGTGTGTGTAGGTGTGATATGGAGTTTATCTTAATATTTAT |
| Delete-JEN2-fw | ATCATACACATATACATATATATAATAACTAACCATAGAAATATTGTGTTGGTGGTATTTTATTTATTTATGTTTAGATTATCATATAA |
| Delete-JEN2-rv | TTATATGATAATCTAAACATAAATAAATAAAATACCACCAACACAATATTTCTATGGTTAGTTATTATATATATGTATATGTGTATGAT |
